# The tortoise strategy as an arbovirus fitness phenotype within the mosquito as revealed by a novel formulation of age-structured vectorial capacity

**DOI:** 10.1101/552125

**Authors:** E. Handly Mayton, A. Ryan Tramonte, Helen J. Wearing, Rebecca C. Christofferson

**Affiliations:** Department of Pathobiological Sciences, School of Veterinary Medicine, Department of Pathobiological Sciences, Baton Rouge, Louisiana; Department of Mathematics, University of New Mexico, Albuquerque, New Mexico; Center for Computation and Technology, Louisiana State University, Baton Rouge, Louisiana

**Keywords:** EIP, vector competence, Aedes, vectorial capacity, Zika, arbovirus, extrinsic incubation period, mortality, biting rate

## Abstract

The transmission dynamics of arboviruses like Zika virus (ZIKV) are most often evaluated by vector competence and the related extrinsic incubation period (EIP), which represent the proportion of vectors that become infectious given exposure and the time it takes for a vector to become infectious given exposure, respectively. Thus, EIP is the temporality of vector competence, and these measures have been used to evaluate the relative fitness of arbovirus systems. However, another temporal process critical to assessing arbovirus transmission dynamics is the age-structure of vector populations, as studies have demonstrated how vector mortality interplays with vector competence and EIP to alter transmission system efficiency. These and other parameters are critical to vectorial capacity (VC), a measure of transmission potential of a vector-pathogen system. However, how these three components – EIP, vector competence, and age – affect VC still needs to be addressed. We first compared experimentally how vector competence/EIP and mosquito age at the time of infection acquisition (Age_acquisition_) interacted in an *Aedes aegypti-*ZIKV model system. We found that Age_acquisition_ did not alter the vector competence/EIP using traditional analyses, except in the context of mortality. To capture and quantify this age-dependent context, we developed an age-structured vectorial capacity framework (VC_age_) by experimentally determining daily mortality and probability of biting, as well as vector competence/EIP parameterized as EIP_Min_ and EIP_Max_. Like previous studies, we found that arbovirus phenotypes leading to outbreaks are not straightforward and may follow a tortoise and the hare (TotH), whereby slow and steady is as or better than fast and furious phenotypes. Understanding the contributions of these age-dependent life traits as well as VC_age_ allows for quantification and visualization of both the magnitude and temporality of transmission dynamics in an age-dependent manner, which reveals this TotH model that should change how compare and rank arbovirus phenotypes, and perhaps even how we identify ‘highly’ or ‘negligibly’ competent vectors.

## Introduction

The transmission dynamics of arboviruses such as Zika virus (ZIKV) are evaluated over several characteristics, namely vector competence and the extrinsic incubation period (EIP). Vector competence is the ability of a mosquito to acquire and ultimately transmit a virus [1, 2]. The time it takes for this process to occur is referred to as the extrinsic incubation period (EIP) [3]. Vector competence and EIP are interrelated measures of the proportion of vectors that become infectious given exposure and the time it takes for a vector to become infectious given exposure, respectively. Thus, EIP can be described as the temporality of vector competence, and this measure has been used to evaluate the relative fitness of arbovirus systems, which can be affected by extrinsic and intrinsic factors.

Indeed, changes in arboviruses fitness and thus transmission dynamics have been based on altered vector competence, especially as a critical component of vectorial capacity [2, 4-12]. The composite measure of vector competence and EIP into a single, dynamic measure allows for a more comprehensive understanding of this process [1, 13-16]. Not all mosquitoes that are exposed will be able to transmit (vector competence) and the time it takes for those mosquitoes that will transmit is not a constant (EIP), and so understanding this composite over several days post infection is critical [2, 17].

Vectorial capacity (VC) was derived as a measure of transmission potential of a vector-borne pathogen by a competent vector, and incorporates both vector competence and EIP [2, 3, 5, 6, 13, 18]. VC is the vector-centric analogue to the basic reproduction number (R_0_), which describes the number of secondary infectious bites resulting from the introduction of a single infectious vector. VC is given by:

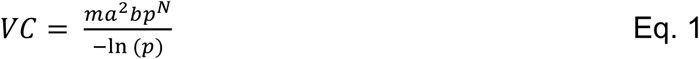

where ‘m’ is the density of mosquitoes relative to humans, ‘a’ is the biting rate, ‘b’ is the vector competence of the mosquito for a particular virus, ‘N” is the extrinsic incubation period, and ‘p’ is the daily probability of mosquito survival [2]. VC, like R_0_, is a threshold parameter, whereby a VC value < 1 is not thought to be able to sustain transmission and a VC value ≥1 is likely to result in an outbreak [19].

VC uses an average daily probability of survival to calculate the probability of a mosquito living through the EIP (p^N^) and the average infectious period (1/-ln(p)). It is intuitive how mortality could impact transmission, but most studies address mortality based on a constant age at which the mosquito acquires the infection [14, 20-23]. The age-dependence of within-vector dynamics is also an important component for determining the relative contribution of EIP and age to VC.

Thus, herein, we experimentally test this hypothesis that transmission dynamics of arboviruses is more affected by time as a function of age versus time as a function of days post-infection. We further develop an age-structured vectorial capacity equation (VC_age_) to quantify these potential effects.

## Materials and Methods

We first wanted to determine if age and/or prior bloodmeals affected the within-mosquito viral dynamics, as well as various life-traits of the mosquito. To that end, we designated three treatment groups: YOUNG, OLDER, and S.OLDER. The YOUNG group was offered an infectious bloodmeal at approximately 5 days post-emergence (dpe). The OLDER group was offered a non-infectious bloodmeal at 5 dpe and then an infectious bloodmeal 1 week later (12 dpe). Further, the S.OLDER group was not offered a prior, non-infectious bloodmeal, but a single infectious bloodmeal at approximately 12 dpe (to match the OLDER timing).

### Virus and mosquitoes

ZIKV strain PRVABC59 (ZIKV-PRV), originally isolated from a human patient in Puerto Rico in 2015, was provided by Dr. Barbara Johnson at the US Centers for Disease Control and Prevention. Prior to use, it was passaged three times in Vero cells and cultured as in [24]. Supernatant was collected 3 days post-inoculation and titer determined as previously described [24]. Titer of ZIKV was verified by qRT-PCR at approximately 8 × 10^7^ plaque forming units (pfu)/mL, matched across all exposure experiments. Virus used for mosquito exposure was never frozen. Colony *Aedes aegypti* (Rockefeller) were provided by Dr. Daniel Swale of the LSU Entomology Department. To isolate the effect(s) of age, mosquitoes were maintained at constant conditions as in [17] with 16:8 light/dark periods and at 28°C constant temperature. Sucrose solution was removed 24 hours before experiments.

### Blood-feeding and oral exposure of *Ae. aegypti*

Bloodmeals were prepared with either ZIKV-infected cell culture supernatant or non-infected cell culture supernatant. Whole bovine blood in Alsever’s solution from Hemostat Labs (Dixon, CA) was used in a 2:1 blood to supernatant ratio [17]. Mosquitoes were fed via the Hemotek (Discovery Labs, UK) membrane feeding system for 45 minutes, after which mosquitoes were cold anesthetized and blood-fed females were placed into clean cartons until further treatment. All groups were starved 24 hours prior to days 5 and 12 post-emergence (regardless if that group received a bloodmeal) and all groups were cold anesthetized and moved to a new carton at days 5 and 12 post-emergence (again, regardless of whether they got a bloodmeal). Thus, all cohorts were treated in the exact same way with the exception of treatment conditions. The experimental design is depicted in Supplementary Fig. S1.

### Within-mosquito kinetics

We wanted to 1) determine if timing of infectious blood-meal affected the within-mosquito viral kinetics and 2) describe the EIP of the virus over the lifetime of the mosquito [3, 25]. To accomplish 1) mosquitoes were sampled at 5, 8, and 11 days post-infection (dpi) to test for infection and dissemination across all groups. Mosquito legs and bodies were put into separate tubes containing 900 μL BA-1 diluent media and BBs. Samples were then homogenized twice at 25 Hz for 3 minutes using a Qiagen Tissuelyzer. RNA was extracted and tested for the presence of viral RNA as in [26]. Each treatment was repeated a total of three times and data are averages over these replicates. Differences among the treatment groups for infection and dissemination rates were tested by a chi-square test for multiple proportions on a day-by-day basis.

### Transmission Assay

To directly assess transmission potential, 20 mosquitoes per treatment were force-salivated. Briefly, ZIKV-exposed mosquitoes were immobilized on ice before removing legs and wings. Mosquitoes were then placed on double-sided tape, and the proboscis of each mosquito was placed into a pipette tip containing 35 μL FBS with 3 mmol/L ATP for 30 minutes, as previously described in [27]. Contents of the pipette tip were ejected into 100 μL BA-1 diluent and stored before testing (see below). Mosquitoes from the two OLDER groups were sampled at 5, 8, and 11 dpi, as well as 16 dpi to represent the end of the mortality study (28 dpe). To accomplish 2), mosquitoes in group YOUNG were sampled at 5, 8, and 11 dpi, as well as additional days post infection in order to match the age of the older groups (12, 15, 18, and 23 dpi). Samples were tested for the presence of viral RNA in saliva using the techniques described above [26, 27]. Differences among the treatment groups transmission rates were tested by a chi-square test for multiple proportions on a day-by-day basis.

### Mortality Study

Mortality studies were performed for the same three treatments (YOUNG, OLDER, and S.OLDER). We added additional mock bloodmeal controls (that is, accounting for any infectious bloodmeal-associated alteration of mortality) where a mock bloodmeal was used in place of infectious bloodmeals. The three controls were: 1) a mock bloodmeal at 5 dpe (M.Y) to correspond to the YOUNG treatment, 2) a mock bloodmeal at 5 dpe, followed by another mock bloodmeal at 12 dpe (M.M) to correspond to the OLDER two bloodmeal treatment, and 3) a mock bloodmeal at 12 dpe (S.M) to correspond to the S.OLDER, one bloodmeal treatment. An additional negative control treatment was performed where the mosquitoes were never blood fed (S). All treatments were cold anesthetized at 5 and 12 dpe, regardless of whether they were offered a bloodmeal so that all mosquitoes experienced the same treatment, and mosquito density per carton was kept relatively constant with an average of 47 mosquitoes/carton (range 36-58). Each mortality treatment was repeated a total of three times and data are averages of the three replicates.

Cartons were checked daily and, when present, dead mosquitoes were counted and removed up to 28 dpe (approximately 1 month), as this has been shown to be the upper limit of believed survival of *Ae. aegypti* and is similar to the range used in Tesla, et al. [27, 28]. Only mosquitoes that took all offered bloodmeals were included in the mortality analyses.

To test for differences in mortality rates among treatments, Kaplan-Meier survival analyses were conducted and the average time to death (TTD) estimated. Daily mortality estimates relative to age were then predicted using best fits in R.

### Determining age-structured willingness to probe

One of the most influential parameters for determining transmission dynamics of vector-borne pathogens is the biting rate. Biting rate is sometimes parameterized as the reciprocal of the number of days between feeds [29]. This assumes that the waiting time between bites takes an exponential form, and we make this assumption for four biting rates: .5 (once every two days), 1 (once a day), and 2 (twice a day) [30]. However, we wanted to determine if the willingness of the mosquito to probe was affected by 1) timing of infectious bloodmeal and/or 2) age of the mosquito. While mosquito biting is a function of many factors, it has been shown that heat cues are sufficient to initiate host seeking behaviors [31]. We use this to determine the willingness or probability of a mosquito probing, which can lead to transmission [32-34].

Ten to twelve mosquitoes per each ZIKV-infected treatment (YOUNG, OLDER, and S.OLDER) were placed in individual, clear plastic canisters (Bioquip) twenty-four hours before being provided a bloodmeal via membrane feeder using 1 mL discs with 800 μL of blood (Hemotek, Discovery Labs, UK). This was done at the same dpi schedule as the vector competence studies above. Willingness to bite was assessed using a two-tiered approach by a single observer to control for observation bias. First, mosquitoes were observed through the clear canister for their general position in the canister and second, the disc was removed to determine if they were on or near the mesh at the top of the canister. In all cases, these two methods of observation matched. That is, if a mosquito was observed to be at the bloodmeal prior to disc removal (looking through the canister), she did not move to the bottom of the canister upon disc removal.

This observation was done at 1, 20, and 45 minutes post placement of the disc and the disc was replaced between observation time points. Thus, a mosquito was assessed as “landed” and recorded as “1” if the female was at the top of the canister at any of the observation times. She was otherwise classified as “not landed” and coded as “0” if she was at the bottom of the canister for all three observation times. We then calculated the probability of biting as a function of age. (Z(t_age_)) was determined by fitting the proportion of mosquitoes that landed or fed at least once a day using a self-starting non-linear least squares regression as above.

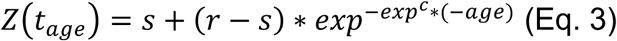

Comparison of probing differences among treatments was assessed both per day post infection as well as per mosquito age by Kruskal-Wallis test.

### Age-structured vectorial capacity

We re-formulated the vectorial capacity equation to estimate VC as a function of the age at which the mosquito acquires an infectious bloodmeal, redefining the parameters with respect to age at the time of acquisition of infection. We define Age_acquisition_ as the age at which a mosquito acquires an infection (the day she takes the infectious bloodmeal) and Age_transmission_ as the age at which she subsequently transmits (Figure 1):

**Figure 1.**
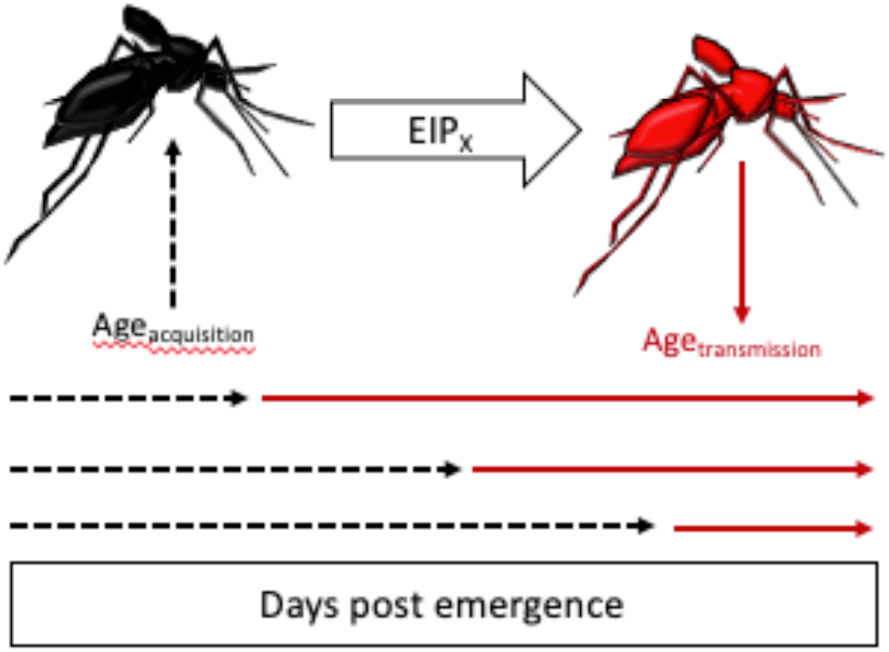
Vector age at time of infection acquisition determines transmission opportunity. Mosquitoes acquire an infection at Age_acquisition_ (black mosquito, left) and after a certain EIP, X% will become infectious (red mosquito, right). The Age_acquisition_ (represented by the black dashed arrows), the opportunity for transmission is altered in that the mosquito has longer or shorter window of transmission before likely time to death (red, solid arrows).

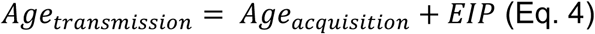

In VC_age_, m is still the mosquito-to-human density and, for illustrative purposes, held constant here at 1. The parameters z_acquisition_ and z_transmission_ are the probability of a mosquito biting at age of acquisition and age at time of transmission, respectively. The traditional calculation of p^N^ represents the probability of a mosquito living through number of days N, the EIP, and p^N^/(-ln(p)) represents the expected infectious days given N and p [35]. In the context of an age-dependent vectorial capacity framework, we can calculate a more precise probability of surviving based on the day the mosquito obtained an infection and the cumulative survival probability to Age_transmission_ where the cumulative probability of living through the EIP given Age_acquisition_ is given by:

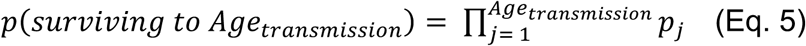

We further estimate the infectious period (L) in an age-structured way by numerically deriving L:

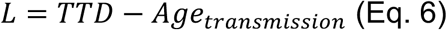

where TTD is the median time-to-death derived from the experimental mortality study and age_transmission_, derived in Eq. 4.

Using the data from our experimental studies, we calculated Age_transmission_ for two scenarios: EIP_min_ and EIP_max_ from our observed transmission data [36], and calculated VC_age_ as:

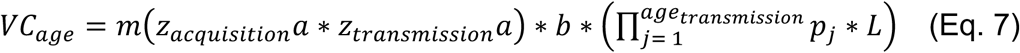

All statistics and subsequent graphics were performed and generated using R version 3.4.3. All functions and function parameters used to fit the data and obtain age-dependent distributions of these parameters are given in Supplementary Table S1. Goodness of fit was assessed either through AIC (for non-linear models) or R^2^ for linear models.

## Results

### Within-mosquito dynamics are more affected by time as a function of age versus days-post-infection

We first wanted to determine if the age at which a mosquito is offered an infectious bloodmeal and acquires the ZIKV infection (Age_acquisition_) impacted the within-vector kinetics of the mosquito. To do this, we looked at infection, dissemination, and transmission of the three treatments in the context of days post-infection. We found no significant difference in the infection or dissemination rates across all treatments (Supplemental Information Table S2). However, when direct age comparisons are made, the effect of Age_acquisition_ becomes obvious an infection (Figure 2).

**Figure 2:**
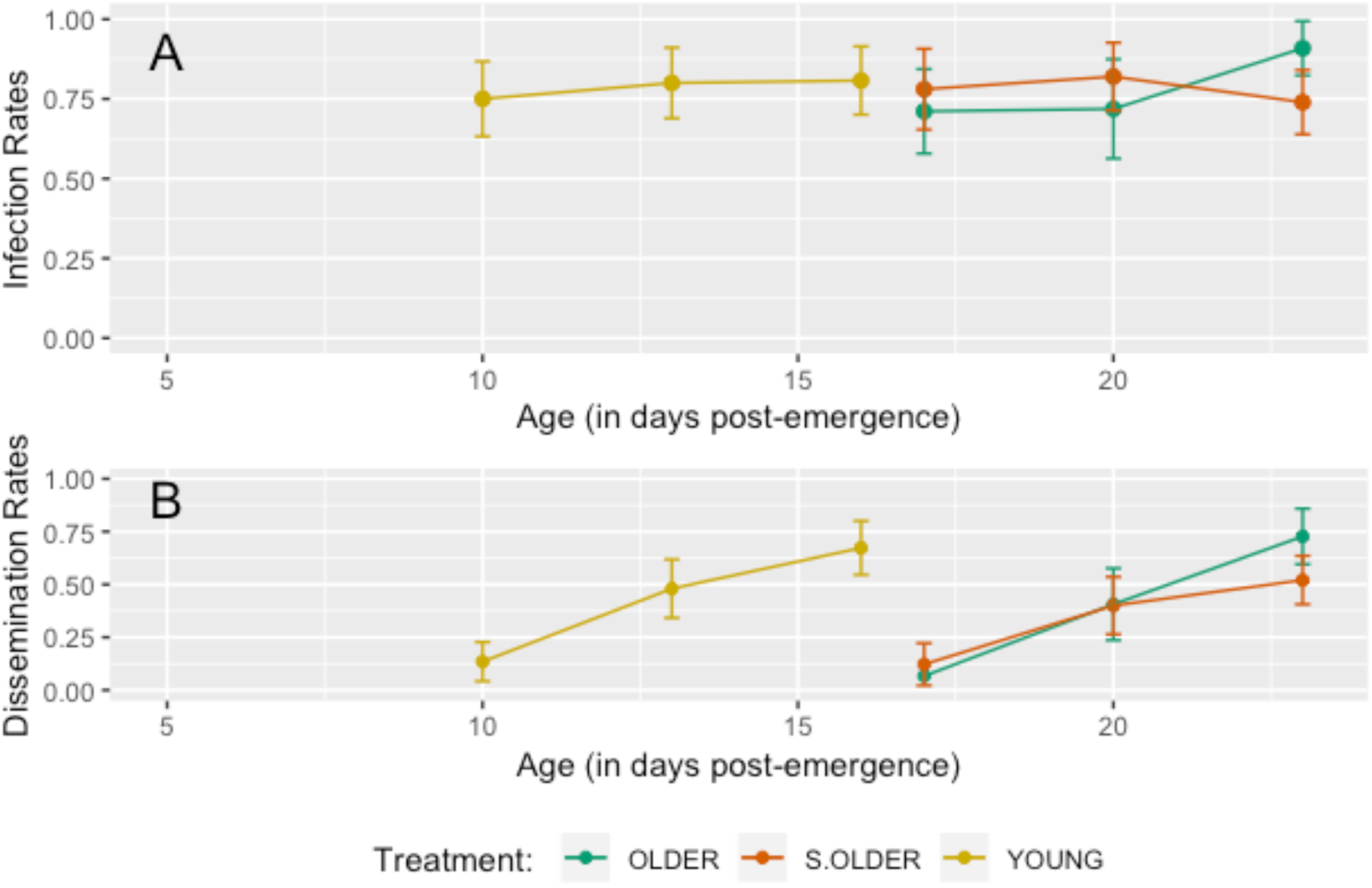
Infection and dissemination rates in the context of mosquito age. Despite no significant effect of treatment on infection and dissemination rates when assessed over days post infection, Age_acquisition_ has an obvious impact on the timing of these processes.

When we evaluated the proportion transmitting in terms of days post exposure (5, 8, and 11 only) none of the treatments were transmitting by on days 5 or 8 dpi. On day 11, the transmission rates got to 5%, 10%, and 15% in groups OLDER, S.OLDER, and YOUNG, respectively. These differences were not statistically significant using the Chi-square test for proportion (p>0.5).

However, when we investigated mosquitoes from group YOUNG at time points that age-matched OLDER and S.OLDER treatments (Table 1), the YOUNG group achieved a maximum transmission of 45% at 28 days old (23 dpi) versus only 15% for the S.OLDER group and 10% for the OLDER group at 28 days old (16 dpi).

**Table 1:**
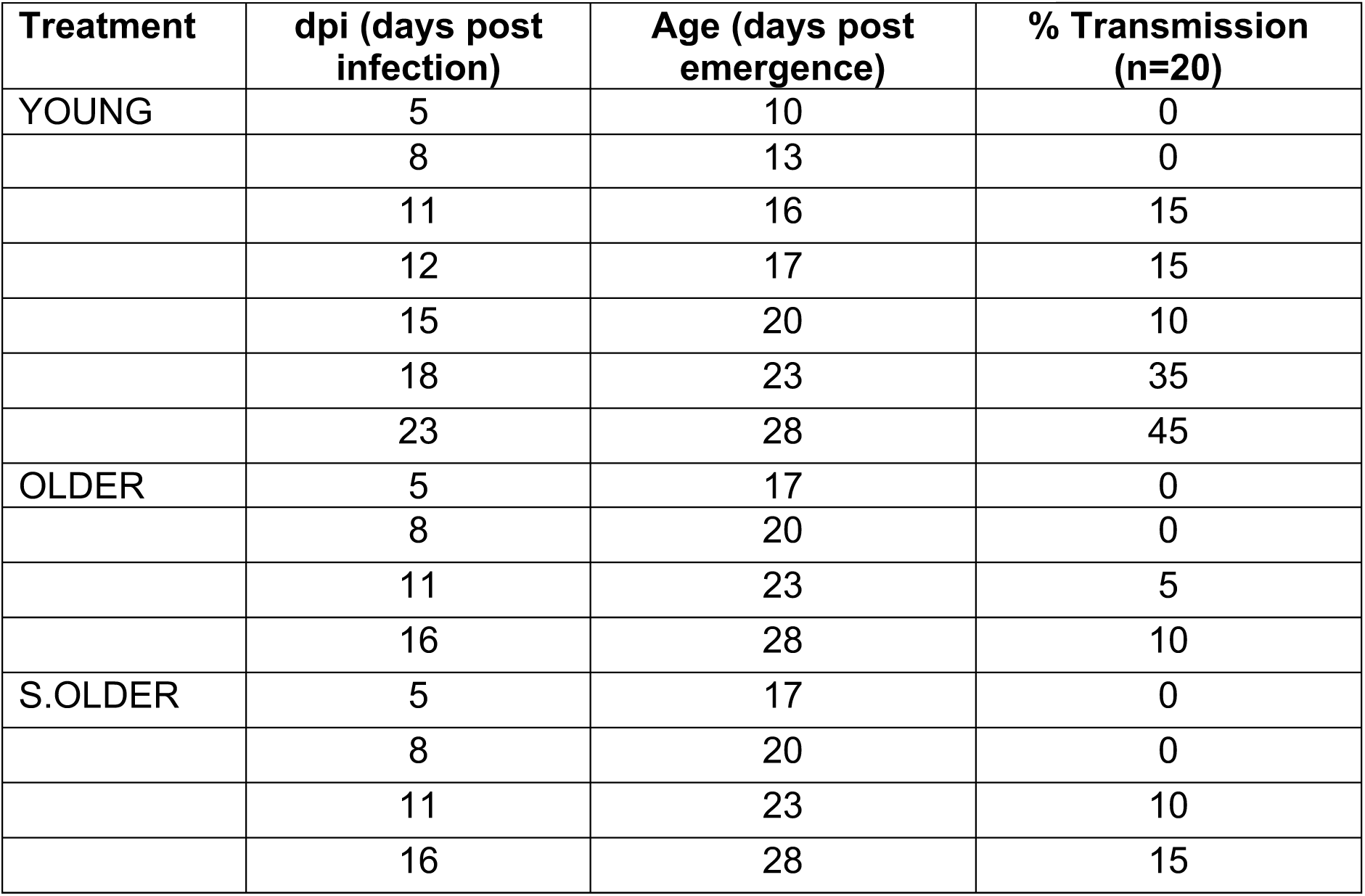
Transmission rates for each day post infection (dpi) and corresponding mosquito age for each of the three treatment groups.

| **Treatment** | **dpi (days post infection)** | **Age (days post emergence)** | **% Transmission (n=20)** |
| --- | --- | --- | --- |
| YOUNG | 5 | 10 | 0 |
|  | 8 | 13 | 0 |
|  | 11 | 16 | 15 |
|  | 12 | 17 | 15 |
|  | 15 | 20 | 10 |
|  | 18 | 23 | 35 |
|  | 23 | 28 | 45 |
| OLDER | 5 | 17 | 0 |
|  | 8 | 20 | 0 |
|  | 11 | 23 | 5 |
|  | 16 | 28 | 10 |
| S.OLDER | 5 | 17 | 0 |
|  | 8 | 20 | 0 |
|  | 11 | 23 | 10 |
|  | 16 | 28 | 15 |

The EIP_max_ we define as the earliest EIP (days post-infection) where the maximum proportion of mosquitoes are transmitting, and we define the EIP_min_ as the earliest EIP where the proportion of mosquitoes transmitting is minimal, but greater than 0. EIP values at different vector competence levels have been used to evaluate the vectorial capacity for malaria [36]. For the YOUNG group, EIP_max_ was 23 dpi at 45%, and EIP_min_ was 11 dpi at 15%. For the OLDER group, EIP_max_ was 16 dpi at 10% and EIP_min_ was 11 dpi at 5%. For the S.OLDER group, EIP_max_ was 16 dpi at 15% and the EIP_min_ was 11 dpi at 10%.

### Mortality of *Aedes aegypti* is modestly affected by timing of infectious bloodmeal

When each infectious treatment was compared to a matching non-infectious control group, the only significant difference in median time to death (TTD) was between the S.OLDER and S.M treatments, with an estimated difference in TTD of two days (Supplementary Table S3). Of interest, the non-blood fed sugar-only controls died significantly faster than any of the blood-fed treatments with an average TTD of 19.6 days (Supplementary Table S3), which has been previously shown [37].

Pairwise comparisons of the ZIKV-exposed treatments determined that group YOUNG had a significantly longer median time to death (TTD) when compared to groups OLDER and S.OLDER, though this difference was modest (0.6 and 1.4 days, respectively) (Figure 3). The TTD for the YOUNG group was 25.9 days, 25.3 days for the OLDER group, and 24.5 days for the S.OLDER group. This corresponds to average daily survival probabilities of 0.962, 0.961, and 0.960 for the YOUNG, OLDER, and S.OLDER groups, respectively Predicted daily survival rates were generated for the YOUNG group using a non-linear fit and a linear model was fit to the OLDER and S.OLDER groups. The parameters of these models and goodness of fit assessments are given in Supplementary Table S1, and the observed and predicted values are shown in Supplementary Figure S2 for all three treatment groups. Additional comparisons were made among the treatment and control groups, detailed in Supplementary Supporting Text S1, Supplementary Figure S3, and Supplementary Table S3.

**Figure 3.**
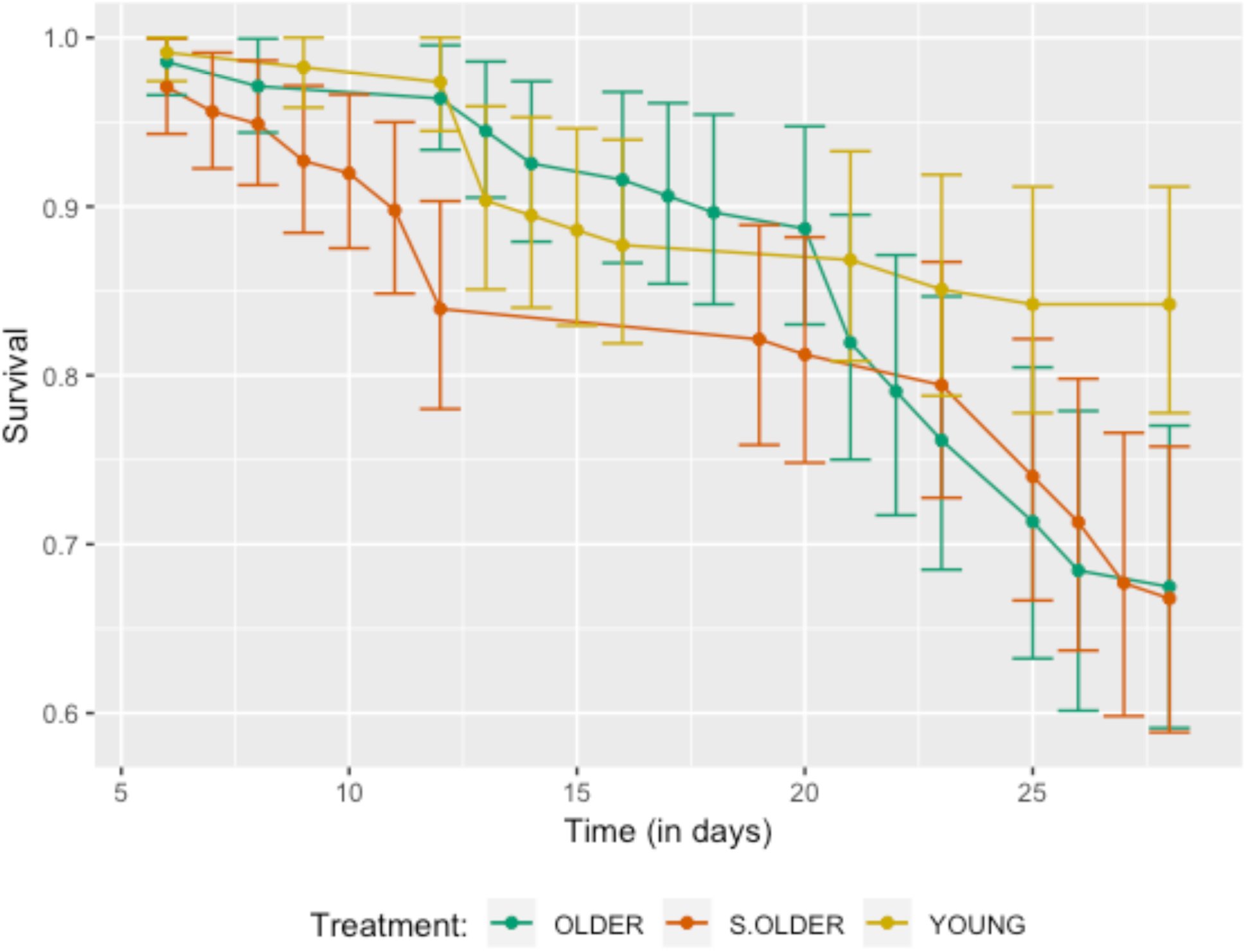
Survival curves of female *Ae. aegypti* by treatment. Each line represents the combined data from three replicates per treatment: YOUNG (gold line), OLDER (green line), and sugar-OLDER (S.OLDER, red line). Average time to death of treatment YOUNG was significantly, but modestly longer than treatments OLDER and S.OLDER.

### Age-dependence of willingness to probe

There was no significant difference in the willingness to probe based on treatment (YOUNG, OLDER, S.OLDER) at each day post infection via Kruskal-Wallis test, but there was a significant effect of age (p<.05). We then were able to fit a daily probability of probing based on age (Figure 4).

**Figure 4.**
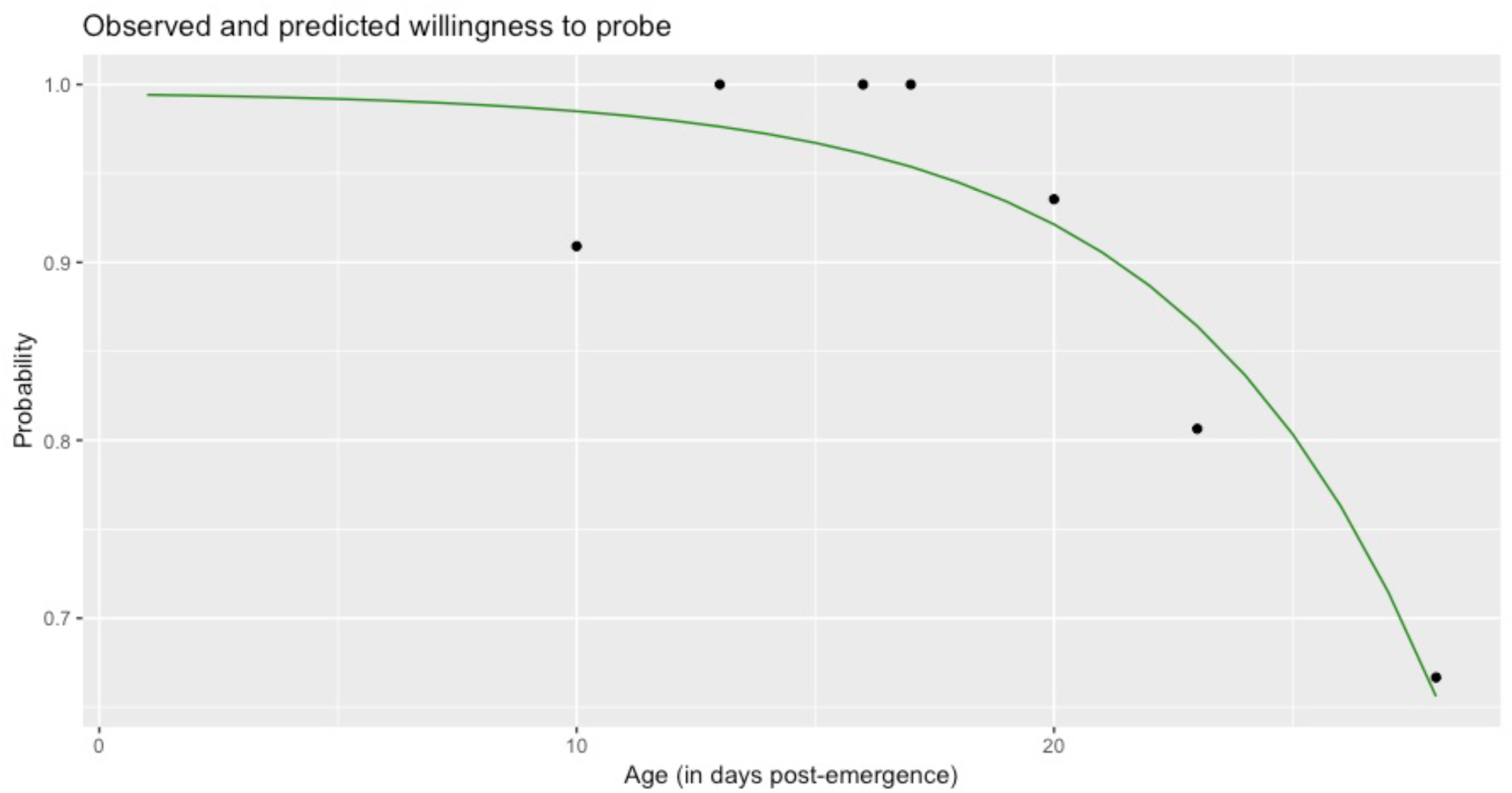
Observed and predicted probabilities of daily biting. The observed daily biting frequencies (dots) from the laboratory experiments and the fitted daily predictions (green curve).

### VC_age_ framework for assessing transmission as a function of age

To calculate VC_age_, we used treatment-group specific median TTD and predicted daily probabilities of survival (Supplemental Figures 4) and the overall daily prediction of willingness to bite in Figure 4. Since biting rate (the number of bites per day) is a field-derived parameter, we calculated VC_age_ over three biting rates of 0.5, 1, and 2; and the mosquito-to-human density was held constant at m=1 [17]. For comparison, we calculated the traditional VC using the average life-dependent traits determined experimentally above, EIP_min_ and EIP_max_, and the biting rates above. These are given in Table 2.

**Table 2:** Traditional VC calculations for EIP_min_ and EIP_max_ for each treatment group (calculated according to Eq. 1).

| Treatment | EIP | Bite rate | VC |
| --- | --- | --- | --- |
| YOUNG | $EIP_{max}$ | 2 | 19.06 |
|  |  | 1 | 4.77 |
|  |  | 0.5 | 1.19 |
| | $EIP_{min}$ | 2 | 10.11 |
|  |  | 1 | 2.53 |
|  |  | 0.5 | 0.632 |
| OLDER | $EIP_{max}$ | 2 | 4.03 |
|  |  | 1 | 1.01 |
|  |  | 0.5 | 0.25 |
| | $EIP_{min}$ | 2 | 6.49 |
|  |  | 1 | 1.62 |
|  |  | 0.5 | 0.41 |
| S.OLDER | $EIP_{max}$ | 2 | 7.65 |
|  |  | 1 | 1.91 |
|  |  | 0.5 | 0.48 |
| | $EIP_{min}$ | 2 | 6.25 |
|  |  | 1 | 1.56 |
|  |  | 0.5 | 0.39 |

### YOUNG GROUP

While traditional VC calculations indicated that there was outbreak potential at all biting rates for this group, this was not the case when VC_age_ was used. At EIP_max_, the VC_age_ at biting rates of 0.5 (once every two days) or 1 (once a day) indicates no likelihood of outbreak (VC_age_<1). The window of opportunity, which encompasses the Age_acquistion_ that results in VC_age_ greater than 1, meaning this is the necessary window of time a mosquito must acquire an infection in order to spark an outbreak. The window of opportunity for EIP_max_ was only 2 days at a bite rate of 2 and the range of VC_age_ 1.40-3.01 (Figure 5). For EIP_min_, again a bite rate of 0.5 did not result in a VC_age_ greater than 1. The window of opportunity was 12 days for a bite rate of 2 and 7 days for a bite rate of 1. The range of VC_age_ was 1.19-7.90 for bite rate of 2 and 1.09-1.98 for a bite rate of 1 (Figure 5).

**Figure 5:**
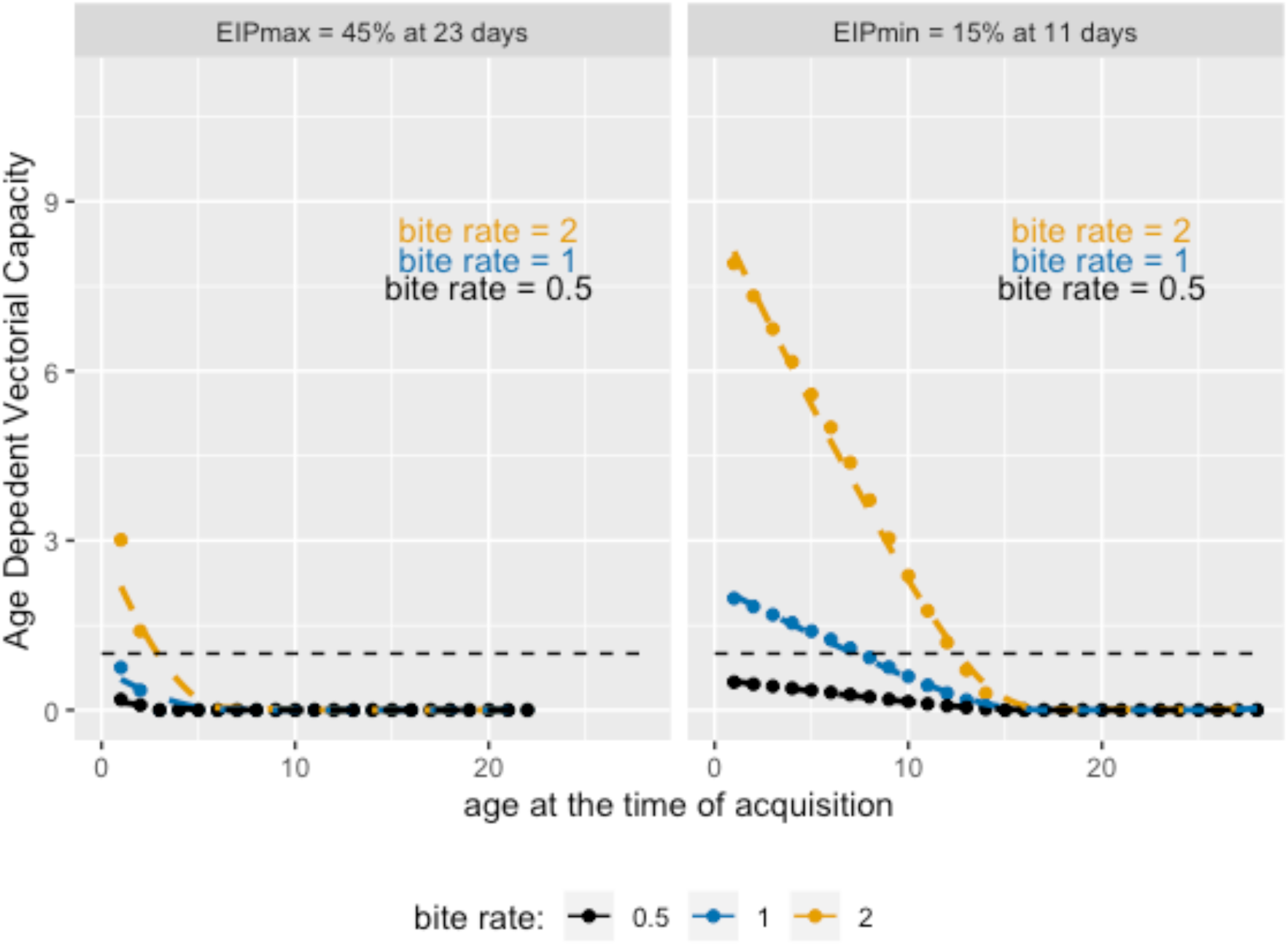
Vectorial capacity calculations using VC_age_. The age-dependent vectorial capacity values (y-axis) depending on the age at acquisition (x-axis) depending on bite rates of 2 (yellow line), 1 (blue line), and 0.5 (black line) for (**LEFT**) EIP_max_ of the YOUNG group, = 45% at 23 dpi, and **RIGHT**) EIP_min_ of the YOUNG group = 15% at 11 dpi.

### OLDER and S.OLDER groups

For both the OLDER and S.OLDER groups, the infectious bloodmeal occurred at 12 days old and at that time point and beyond, VC_age_ < 1 for all parameter combinations (Supplemental Figure S3). This means that Age_acquisition_ of 12 days is too late given these particular characteristics (biting willingness, mortality, and EIP). With these vector age-dependent traits, both EIP and vector competence would need to be altered significantly to achieve a VC_age_ ≥ 1 at 12 days of age (when exposure occurred).

For example, we consider a hypothetical EIP_max_ of EIP_50_ (the time it takes for 50% of exposed mosquitoes to transmit) to demonstrate how these group-specific mortality rates and the age-structure of willingness to bite interact to drive vectorial capacity in the context of Age_acquisition_ [28, 38]. The S.OLDER group would require an EIP_50_ of 11 dpi or 9 dpi for biting rates of 2 and 1, respectively. For the OLDER group, the EIP_50_ would need to be 12 dpi or 10 dpi for biting rates of 2 or 1, respectively, in order for VC_age_ to be greater than 1 at Age_acquisition_ of 12 days. A biting rate of 0.5 was not permissive to VC_age_ ≥ 1 for the S.OLDER group, while an EIP_50_ of just 3 days would theoretically allow for VC_age_ ≥ 1 in the OLDER group, though this short EIP_50_ is biologically unlikely.

Given, though, that the EIP_min_ was also capable of supporting transmission perpetuation, we investigated how fast an EIP would need to be at lower limits of vector competence, utilizing an EIP_10_ scenario [36]. For the OLDER group, an EIP_10_ of 9 days resulted in VC_age_ = 1.34 at 12 dpe for a biting rate of 2, while for the S.OLDER group, an EIP_10_ of 7 days at a biting rate of 2 resulted in a VC_age_ = 1.20 at 12 dpe. These results indicate that an EIP_10_ of 8 or 9 days or less would support transmission at this older Age_acquisition_ (for the S.OLDER and OLDER groups, respectively). For a biting rate of 1 and 0.5, VC_age_ never reached above 1 for 12 dpe for EIP_10_.

## Discussion

Prediction of vector-borne disease spread remains difficult, as transmission of vector-borne disease is a dynamic, multifaceted system. This includes life traits of the mosquito, environmental factors, and vector-virus interactions [4, 11, 39-41]. Here we demonstrate that mosquito age at the time of pathogen acquisition is a powerful driver of transmission potential due, in large part, to the age-dependence of daily mortality and biting habits. Further, these drivers lead to large differences in the quantification of outbreak potential. In this case, it was the difference between declaring a risk for outbreak or not (Figure 5). Many things affect vector competence including vector species, discrete populations within species, and environmental factors [1, 14, 19, 20, 28, 40, 42, 43]. Several recent studies have focused on environmental factors such as temperature, and found that temperature not only affects vector competence of many arboviruses, especially in *Aedes aegypti*, but also several life traits of the mosquito [27, 39, 41]. This means that transmission is ultimately affected by the interactions of all of these temporally dependent processes, and while mathematical frameworks for assessing temperature and other extrinsic factors have been developed, they do not often explicitly consider age, another important temporal component to the transmission cycle of vector-borne diseases.

Our treatments, which focused upon the age at the time of infectious bloodmeal, showed no significant impacts on the vector competence of colony *Ae. aegypti* for ZIKV. One such study has shown the effects of multiple bloodmeals [44]. While one of our treatments did include multiple bloodmeals, this study was not focused upon the effects of multiple bloodmeals on the transmission of ZIKV by *Ae. aegypti*, and our study, delivered the second bloodmeal at a much longer interval (7 days versus 3 dpi in [44]) which likely plays a role in this disparate finding and indicates that there is likely age-dependence in this phenomenon, as well. However, this hypothesis is outside the scope of the current study.

Further, we held the density constant. For our study and in the context of vectorial capacity, this is appropriate since VC is looking at the kick-off event of an outbreak and can be evaluated based on just one mosquito becoming secondarily infected. As this parameter is scalar, increasing it would have a linear and thus relatively proportional effect on VC_age_. The technology for determining the age-structure of natural mosquito populations in the field is currently still in development. For example, a study using near-infrared spectroscopy was able to predict the age of female *Ae. aegypti* +/-2 days, indicating that determining the age-structure of a mosquito population is possible, and that such technology could be refined for field studies [45, 46]. Further, mid-infrared spectroscopy had varying, but some promising results in determining the age-structure of *Anopheles* mosquitoes [46, 47]. As these technologies are pursued and refined, there will be a need for ways to understand and quantify age-dependent interactions among vector competence, EIP, mosquito lifespan, and biting behavior [4].

Finally, traditional thinking often prioritizes higher vector competence as a penultimate measure of fitness, as it has been used to comparatively rank viral fitness [2, 4, 8-12, 32]. But our results indicate, especially at older Age_acquisition_, that a shorter EIP is more critical than the magnitude of vector competence. Similarly, a study by Althouse et. al. also found that the temporality of transmission from non-human primates was sometimes more impactful than the magnitude of the viremia leading to transmission to the mosquito [48]. They proposed a “tortoise-or-the-hare” (TotH) model to describe this relationship between arboviral viremia profiles in non-human primates and the predicted transmission success to vectors, showing that the strategy of “slow and steady” viremia – lower levels for longer periods–resulted in higher predicted transmission success of arboviruses [48]. Here again we invoke the TotH model to describe the complimentary side of the vector-borne pathogen transmission cycles, in the direction of vector-to-vertebrate. Overall, we demonstrate that (provided the biting rate is high enough) EIP_min_ results in a longer window of opportunity compared to EIP_max_ and is thus at least as, if not more capable of producing an outbreak than the EIP_max_ scenario, though the magnitude of the initial outbreak may be less. However, this same TotH model recently described macro-transmission dynamics in Colombia, where it was demonstrated that slow burn-in epidemics actually resulted in cumulatively more cases and higher R_0_ values than in initially explosive outbreaks [49]. Thus, the EIP_min_ scenario may lead, at a macro-level, to more prolonged transmission as it would results in a slower burn-in than the EIP_max_ scenario.

The TotH metaphor is applicable to the micro-level transmission dynamics both within the vertebrate and within the vector, as well as to the macro-level transmission dynamics of affected human populations. Understanding the contributions of these age-dependent life traits as well as VC_age_ allows for quantification and visualization of both the magnitude and temporality of transmission dynamics in an age-dependent manner, which reveals this TotH model that should change how compare and rank arbovirus phenotypes, and perhaps even how we identify ‘highly’ or ‘negligibly’ competent vectors.

## Acknowledgements

We would like to thank all our lab mates for putting up with the hogging of freezer space to hold all of the samples generated for this study. And last but not least, we’d like to thank Aesop for his apparently ubiquitously applicable fable.

## Bonus log line

So perhaps instead of hunting wabbits, we should be looking out for the tortoise.

