## Supplemental Information for "The tortoise strategy as an arbovirus fitness phenotype within the mosquito as revealed by a novel formulation of age-structured vectorial capacity"

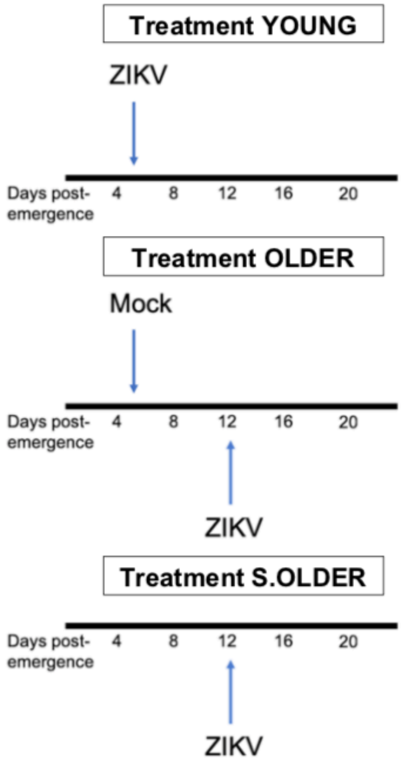


**Supplementary Figure S1. Illustration of main treatment design for vector competence experimentation.** Three treatments were applied to assess our hypothesis. Treatment YOUNG (ZIKV) received an infectious bloodmeal at 5 days old; Treatment OLDER (Mock-ZIKV) received a mock infectious bloodmeal at 5 days old, followed by an infectious bloodmeal at 12 days old; and Treatment S.OLDER (sugar-ZIKV) received an infectious bloodmeal at 12 dpe.

**Supplementary Table S1. Modeled fits of parameters, the type of model, and the parameter values.** For each parameter where predictions were made from experimental data, the type of model (and R self-starting function when applicable) and parameter estimates for each fit are given below. Goodness of fit was assessed either through AIC (for non-linear models) or $R^2$ for linear models.

| Parameter | Model (R self-start function) | Parameter estimates | Goodness of fit |
|---|---|---|---|
| Daily probability of survival for YOUNG group | Asymptotic regression (SSasymp) | s = 0.796<br>r = 1.13<br>c = -2.56 | AIC = -43.8 |
| Daily probability of survival for OLDER group | Linear model | slope = -0.016<br>intercept = 1.13 | Adj. $R^2$ = .88 |
| Daily probability of survival for S.OLDER group | Linear model | slope = -0.012<br>intercept = 1.04 | Adj. $R^2$ = .95 |
| Probability of daily biting | Asymptotic regression (SSasymp) | s = .996<br>r = .995<br>c = -1.665 | AIC = -0.71 |

**Supplementary Table S2. Infection and dissemination rates for each day post-infection (dpi) and corresponding mosquito age for each of the three ZIKV treatments.** Percent infection was determined by the proportion of infected abdomens over total exposed; and percent dissemination was determined as the proportion of infected legs over total exposed. Three replicates were performed for each treatment, but proportions and sample sizes (n) are combined from all three replicates.

| **Treatment** | **dpi (Age)** | **% Infected (n)** | **% Disseminated (n)** |
|---|---|---|---|
| YOUNG | 5 (10) | 77.45 (52) | 38.53 (52) |
| | 8 (13) | 81.90 (50) | 48.25 (50) |
| | 11 (16) | 83.38 (52) | 69.93 (52) |
| OLDER | 5 (17) | 74.67 (45) | 4.00 (45) |
| | 8 (20) | 69.40 (32) | 37.50 (32) |
| | 11 (23) | 87.88 (44) | 66.67 (44) |
| S.OLDER | 5 (17) | 78.27 (41) | 9.80 (41) |
| | 8 (20) | 80.89 (50) | 35.30 (50) |
| | 11 (23) | 74.09 (73) | 51.94 (73) |

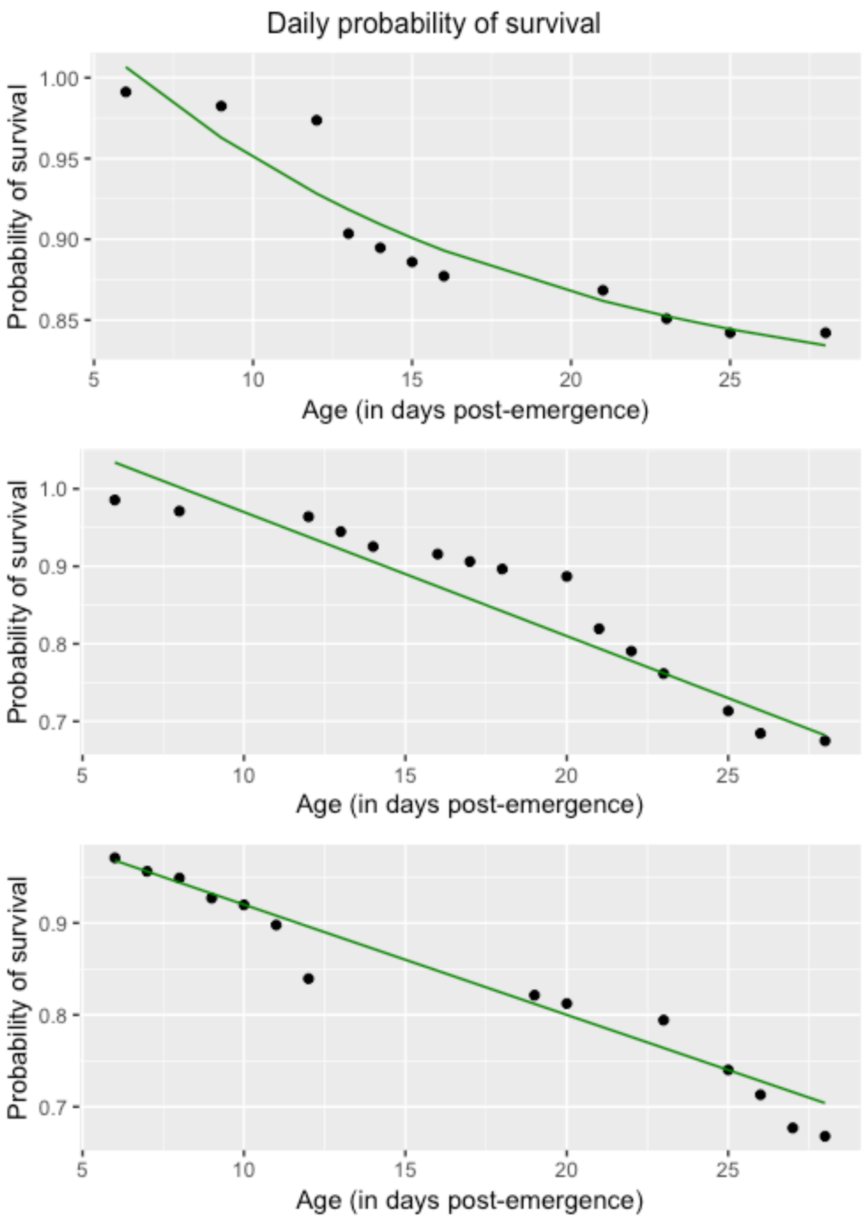


**Supplementary Figure S2. Observed and predicted daily probabilities of survival for the three treatment groups**. Observed daily survival rates (dots) and the predicted daily survival rates (green curve) for YOUNG group (Top), OLDER group (Middle), and S.OLDER group (Bottom).

**Supporting Text S1: Mortality of *Aedes aegypti* with respect to bloodmeals**
The supporting controls for each of the treatment groups were as follows:

1. For YOUNG group: a mock bloodmeal at 5 days post emergence (dpe) followed by sugar sustenance ("M.S")
2. For OLDER group: a mock bloodmeal at 5 dpe followed by a second mock bloodmeal at 12 dpe ("M.M")
3. For S.OLDER group: a mock bloodmeal at 12 dpe only ("S.M")
4. A sugar-only control ("S") that received no bloodmeals.

Of interest, the non-blood fed sugar-only controls (group S) died significantly faster than any of the other treatments with an average TTD of 19.6 days (Figure S3). To determine if there was a generalized effect of exposure on mortality, we compared all groups with a ZIKV exposure to those without, excepting the group that received no bloodmeal at all which was removed from the analysis. There was no significant difference between ZIKV-exposed mosquitoes (groups YOUNG, OLDER, and S.OLDER) and the non-exposed groups ($p>.05$).

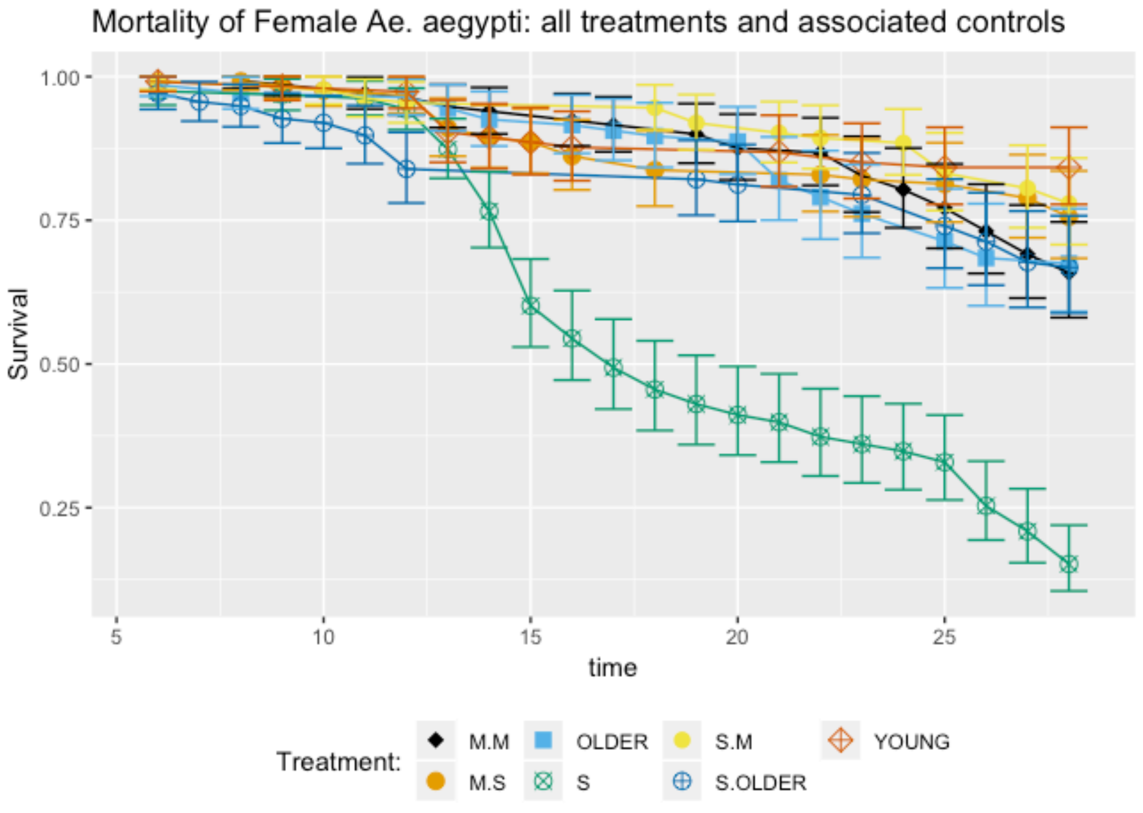


**Supplementary Figure S3:** Mortality curves for all treatments and associated controls. The sugar-only group had significantly faster mortality compared to the other groups which received at least one bloodmeal.

**Supplementary Table S3. Median time to death and sample size for ZIKV-infection treatments and unexposed controls used in the mortality study.** Exposed groups were the three treatments exposed to ZIKV: YOUNG -- ZIKV at 5 days post emergence (dpe); OLDER – mock bloodmeal at 5 dpe and ZIKV bloodmeal at 12 dpe; S.OLDER – only a ZIKV bloodmeal at 12 dpe. Unexposed groups were not exposed to ZIKV but matched for bloodmeal uptake: M.S – mock bloodmeal at 5 dpe; M.M – mock bloodmeals at 5 and 12 dpe; S.M – mock bloodmeal at 12 dpe; and S – no bloodmeal, sugar only. Median times to death and total sample sizes per treatment are given below.

| **Group** | **Treatment** | **Time to death** | **Sample size (n)** |
|---|---|---|---|
| Exposed | YOUNG | 25.9 | 114 |
| | OLDER | 25.3 | 139 |
| | S.OLDER | 24.5 | 137 |
| Unexposed | M.S | 25.5 | 123 |
| | M.M | 25.8 | 139 |
| | S.M | 26.5 | 132 |
| Sugar Only | S | 19.6 | 158 |

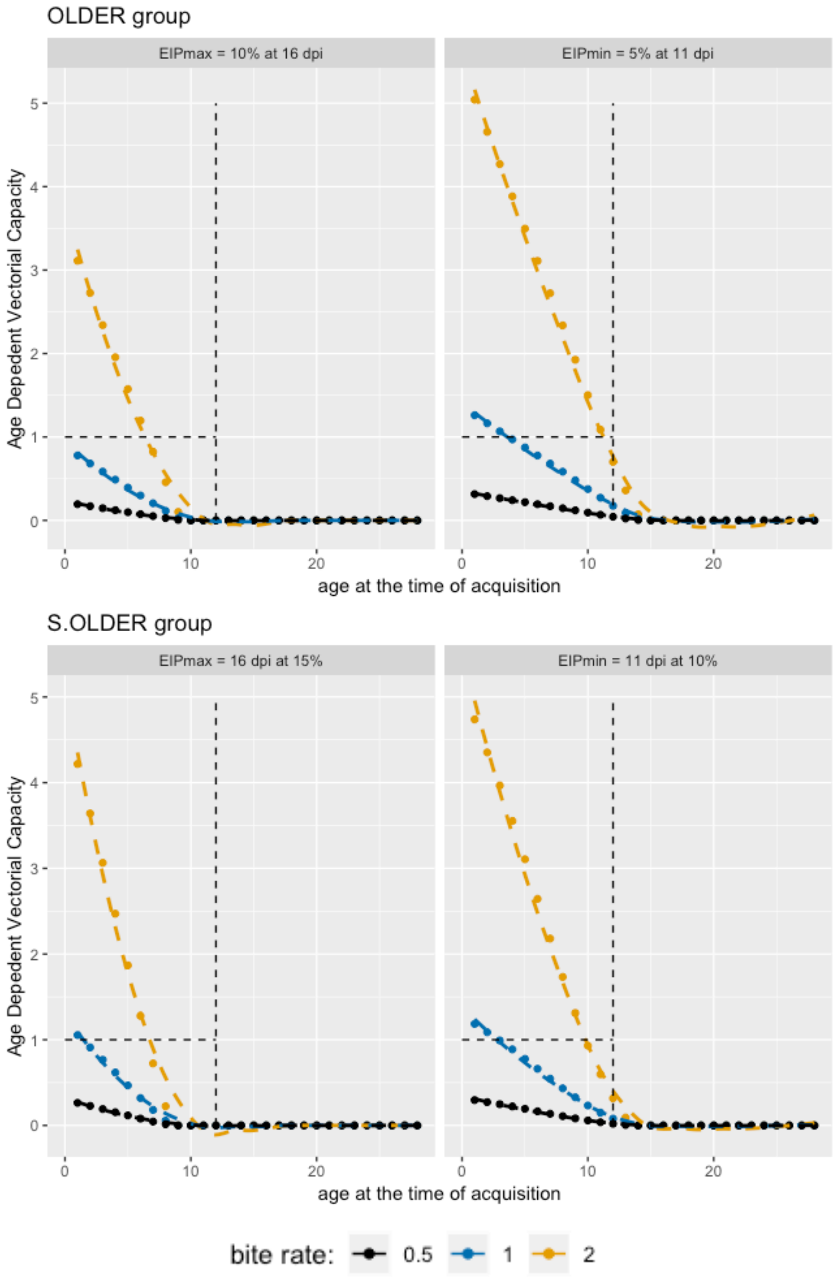


**Supplemental Figure S4**. $VC_{age}$ values for the OLDER and S.OLDER groups indicates that at $Age_{acquisition}$ (12 days post emergence), none of these scenarios resulted in $VC_{age} \geq$ 1. VCage was only greater than 1 when calculated at an $Age_{acquisition}$ was well before the actual 12 dpe. Dotted lines are where $VC_{age}$ = 1 (horizontal) and mosquito age = 12 (vertical)
